# Stability and heterogeneity in the anti-microbiota reactivity of human milk-derived Immunoglobulin A

**DOI:** 10.1101/2023.03.16.532940

**Authors:** Chelseá B. Johnson-Hence, Kathyayini P. Gopalakrishna, Darren Bodkin, Kara E. Coffey, Ansen H.P. Burr, Syed Rahman, Ali T. Rai, Darryl A. Abbott, Yelissa A. Sosa, Justin T. Tometich, Jishnu Das, Timothy W. Hand

## Abstract

Immunoglobulin A (IgA) is secreted into breast milk and is critical to both protecting against enteric pathogens and shaping the infant intestinal microbiota. The efficacy of breast milk-derived maternal IgA (BrmIgA) is dependent upon its specificity, however heterogeneity in BrmIgA binding ability to the infant microbiota is not known. Using a flow cytometric array, we analyzed the reactivity of BrmIgA against bacteria common to the infant microbiota and discovered substantial heterogeneity between all donors, independent of preterm or term delivery. We also observed intra-donor variability in the BrmIgA response to closely related bacterial isolates. Conversely, longitudinal analysis showed that the anti-bacterial BrmIgA reactivity was relatively stable through time, even between sequential infants, indicating that mammary gland IgA responses are durable. Together, our study demonstrates that the anti-bacterial BrmIgA reactivity displays inter-individual heterogeneity but intra-individual stability. These findings have important implications for how breast milk shapes the development of the infant microbiota and protects against Necrotizing Enterocolitis.

**Summary:** We analyze the ability of breast milk-derived Immunoglobulin A (IgA) antibodies to bind the infant intestinal microbiota. We discover that each mother secretes into their breast milk a distinct set of IgA antibodies that are stably maintained over time.

## Introduction

Breast milk is acknowledged by the World Health Organization and American Academy of Pediatrics as the best source of nutrition for infants (Sobti et al., 2002). Breast milk contains multiple bioactive components, including antibodies, that both prevent infection and aid in the proper installation of the infant microbiota (Gopalakrishna and Hand, 2020; Le Doare et al., 2018; Walker and Iyengar, 2015). IgA, IgG and IgM antibodies are all found in breast milk, but IgA is dominant, making up over 90% of the antibody secreted in the mammary gland. One reason for this is that during pregnancy, IgA-producing B cells are compelled to travel from the intestine to the mammary gland, indicating that IgA secreted into milk is an effort to transfer maternal mucosal immunity to the infant (Lindner et al., 2015; Wilson and Butcher, 2004). During B cell production, IgA is often dimerized by the J-chain which promotes binding and transcytosis of IgA by the polymeric glycoprotein Ig receptor (pIgR) that upon secretion remains bound to IgA as ‘Secretory Factor’ (SF) (Hand and Reboldi, 2021). SF-bound of ‘secretory; IgA (SIgA) is protected against proteolytic cleavage in the intestine, substantially increasing the half-life and functionality of IgA half-life and functionality of IgA at mucosal surfaces (Johansen and Kaetzel, 2011). The majority of IgA secreted into breast milk is SIgA (Rogier et al., 2014).

In addition to protecting against infection (Gopalakrishna and Hand, 2020), SIgA is important in shaping the development of the infant microbiota (Planer et al., 2016; Rogier et al., 2014). In breast fed infants, milk is the predominant source of IgA in the first month of life and in mice, lack of maternal IgA affects the development of the microbiota (Gopalakrishna et al., 2019; Mirpuri et al., 2014; Rognum et al., 1992). Infant formula, which lacks all immunoglobulins, is also associated with alterations in the infant microbiota and increased rates of short and long-term diseases (Dixon, 2015; Oddy, 2017). Preterm infants are particularly susceptible to diseases related to improper regulation of colonization by microbiota, like necrotizing enterocolitis (NEC) (Bode, 2018; Cortez et al., 2018; Neu and Walker, 2011; Nino et al., 2016; Warner and Tarr, 2016). The incidence of NEC is significantly increased in formula-fed preterm infants and the promotion of milk feeding in these children has reduced the incidence of this disease (Neu and Walker, 2011; Nino et al., 2016). In a cohort of milk-fed preterm infants, we have demonstrated that in the days directly preceding the development of NEC, there is a substantial reduction in the fraction of intestinal bacteria bound by breast milk-derived IgA, that is not typically observed in infants who do not develop disease (Gopalakrishna et al., 2019). The majority of IgA ‘unbound’ bacteria in preterm NEC infants come from one family: *Enterobacteriaceae,* which has previously been associated with the disease (Gopalakrishna et al., 2019; Pammi et al., 2017). Changes to IgA binding of the infant microbiota could either be caused by a shift in the composition of the microbiota or the anti-bacterial IgA reactivity of the breast milk. The level of heterogeneity in milk-derived IgA between individuals and over time within one mother is not well understood, but due to their intestinal origin(Lindner et al., 2015), the anti-bacterial reactivity of human mammary gland-resident IgA-producing B cells is likely to be highly individualized.

To measure the milk-derived anti-bacterial IgA response we have developed a flow cytometric array that allows us to define the ability of antibodies to bind the surface of different bacterial isolates. Using this array, we have identified significant heterogeneity between different donors in the binding of bacterial isolates by milk-derived IgA. We also observed isolate level variation in IgA binding, within donor samples, to closely related taxa of *E. coli* and other species. In contrast to the inter-individual heterogeneity, hierarchical clustering and Principal Component Analysis (PCA) of longitudinally collected samples showed consistent clustering within donors, indicating that the anti-bacterial IgA reactivity of an individual is stable over the course of one infant. Analysis of milk samples collected over sequential siblings also revealed stability in anti-bacterial IgA reactivity, indicating that the B cells that secrete IgA into breast milk may be maintained long-term. Finally, we demonstrate that Holder pasteurization, which is commonly used to sterilize human donor milk, globally reduces bacterial binding by IgA. Together our data indicates that the anti-bacterial reactivity of milk-derived IgA is heterogeneous between individuals but also surprisingly stable, even over infants separated by years of time. The temporal stability of breast milk-derived IgA reveals a potential weakness of vertical antibody transmission, where maternal antibody responses are uncoupled from infant intestinal bacterial colonization, potentially limiting BrmIgA’s protective effects against infection and NEC (Gopalakrishna et al., 2019).

## Results

### Determining anti-bacterial IgA reactivity of breast milk using a flow cytometric array

Anti-bacterial IgA is predominantly specific to surface antigens and bacterial staining techniques that use bacterial lysates are complicated by irrelevant antibody cross-reactivity against cytoplasmic proteins and nucleic acids (Slack et al., 2009). Therefore, we modified an approach described by Slack and colleagues to measure anti-bacterial IgA specificity of breast milk antibodies by flow cytometry (Moor et al., 2016; Slack et al., 2009). To negate non-specific signals associated with the non-IgA components of breast milk we isolated SIgA via passage over a streptococcal Peptide M column. LDS-PAGE under reducing conditions of the Peptide M bound fraction revealed bands roughly corresponding in size to Secretory Factor (∼80kDa), IgA Heavy Chain (∼60kDa) and Light Chain (∼30kDa) (**Fig. S1A**)(Sandin et al., 2002). J-chain (15kDa) is known to migrate slowly under LDS-PAGE electrophoresis and is the faint band running slightly below the Light chain at ∼25kDa (**Fig. S1A)** (Zikan et al., 1985). We confirmed by Western blot that each of the four components of SIgA were enriched in the Peptide M bound fraction (**Fig. S1B**). To determine the specificity of an IgA enriched breast milk sample for bacterial surface antigens we incubated purified IgA samples on bacterial isolates individually arrayed on a 96 well plate (**Fig. 1A**). Prior to flow cytometric analysis samples were normalized to rough protein content (280nm Absorbance), which corresponds well to the concentration of IgA measured by ELISA (**Fig. S1C**). After incubation with breast milk-derived IgA, bacteria are stained with a mixture of Syto BC and fluorescently-labelled anti-human IgA. Syto BC is a mixture of bacterial cell wall permeable dyes that allow us to discriminate bacteria from similarly sized debris on the flow cytometer (**Fig. 1B**). Syto BC^+^ bacteria can then be assayed for binding by breast milk-derived IgA by assessing the relative fluorescence normalized to a background control of the same bacteria stained only with anti-human IgA secondary antibody (**Fig. 1C**). Analysis of a dilution series of purified IgA samples revealed that the concentration of SIgA used to test bacterial binding (0.1mg/mL) was saturating for the bacteria tested and thus was used as a standard concentration for all further experiments (**Fig. S1D**). To control for non-specific binding of BrmIgA by bacteria we tested a monoclonal antibody specific to HIV against our array and found only very minimal binding, indicating that our array is measuring anti-bacterial IgA responses (**Fig. 1C**)(Yu et al., 2013). Further, binding of BrmIgA to a soil bacterium *Bradyrhizobium japonicum*, demonstrated marginal signal, indicating that the breast milk-derived IgA response is focused on bacterial taxa that commonly colonize humans (**Fig. 1D**)(Haas et al., 2011).

**Figure 1.**
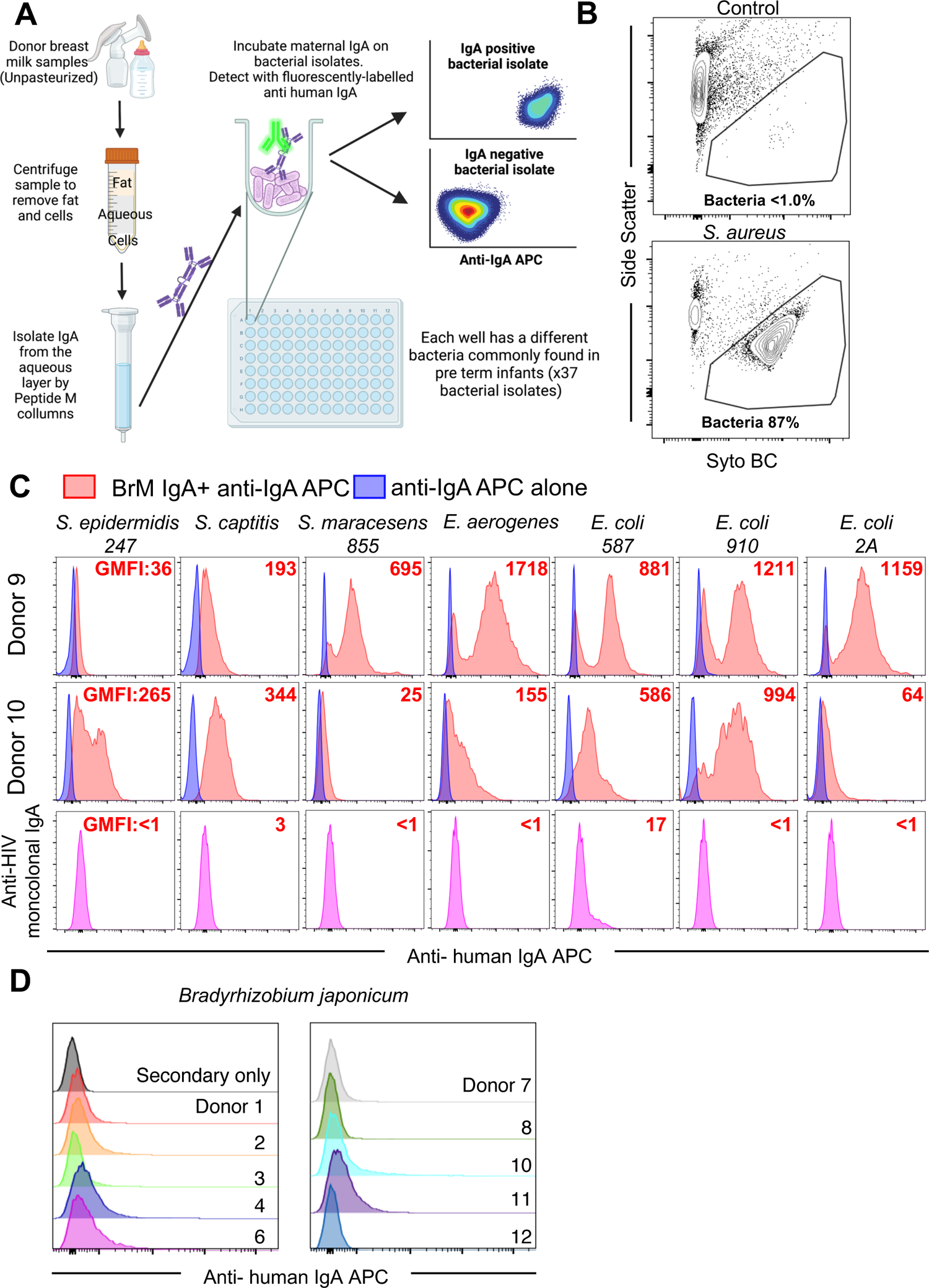
A flow cytometric array for measuring the anti-bacterial specificity of breast milk-derived IgA. **A)** Design of the flow cytometric array. Made with BioRender.com **B)** Examples of SYTO BC^+^/SSC^Dim^ staining used to discriminate bacteria from debris/bubbles in the flow cytometer (control is empty well stained with SYTO BC). Numbers represent the percent of events inside the gate. **C)** Examples of the magnitude of anti-bacterial IgA binding detected in our array comparing two donors (9 and 10) that differ in their anti-bacterial IgA responses. The bottom row shows the reactivity of an anti-HIV IgA antibody against the same bacterial isolates. Numbers in red represent the gMFI of that sample. **D)** Breast milk-derived IgA reactivity, from several donors (as indicated) against the environmental bacteria *Bradyrhizobium japonicum*.

### Heterogeneity in the breast milk-derived anti-bacterial IgA reactivity

After delivery the infant microbiota goes through three main stages (Reyman et al., 2019). First, the infant intestine becomes colonized by common facultative anaerobic bacteria such as *Enterobacteriaceae* and *Enterococcaceae.* Within the next four weeks these bacteria will be supplanted as the dominant taxa by *Bifidobacteria*, which use Human Milk Oligosaccharides as a food source. Six months later, approximately coinciding with the introduction of solid food, there is another switch towards anaerobic *Firmicutes* and *Bacteroidetes* that assist with the digestion of complex carbohydrates. BrmIgA likely contributes to shaping microbiota colonization at all of these stages, but is particularly important for controlling the early bacterial colonizers that comprise the first stage when infants don’t make their own IgA (Gopalakrishna et al., 2019; Koch et al., 2016; Mirpuri et al., 2014; Rognum et al., 1992). Indeed, mouse pups fed by dams that lack IgA production or secretion are colonized with facultative anaerobes such as *Enterobacteriaceae* and *Pastereurellaceae* longer than IgA-secreting controls (Mirpuri et al., 2014; Rogier et al., 2014). Regulation of early colonizing bacteria is especially relevant to preterm infants where increased *Enterobacteriaceae* and in particular IgA-free *Enterobacteriaceae* is associated with the development of Necrotizing Enterocolitis (NEC) (Gopalakrishna et al., 2019; Pammi et al., 2017). Thus, when designing the bacterial array for analyzing the anti-bacterial reactivity of breast milk-derived IgA, we focused on facultative anaerobes, such as *Enterobacteriaceae,* that dominate early infant bacterial colonization. Our array contained 36 individually grown and plated bacterial isolates from 13 different genera that represent the major taxa commonly found in the intestine of preterm infants. All donor samples were normalized for the input concentration of IgA. Analysis of the anti-bacterial IgA responses from 33 donors revealed a substantial amount of heterogeneity, with no two donors being identical (**Fig. 2A**). Thus, individualized differences in the IgA^+^ B cell population of the intestine driven by distinct infection and microbiota experiences likely lead to similar heterogeneity in the breast milk (Bunker et al., 2017; Hapfelmeier et al., 2010; Lindner et al., 2015; Zhang et al., 2017). Comparative analysis of the normalized magnitude of BrmIgA across all bacterial isolates revealed that rather than particular donors making universally strong or weak responses against all bacterial isolates, there was substantial heterogeneity in donor binding from bacterial isolate to isolate (**Fig. 2A-B**). The magnitude of IgA binding to different bacterial isolates was also evenly distributed, where normalized IgA binding for most isolates shared a similar standard deviation (∼1 to 5), though some donors had exceptionally strong binding to different *Staphlyococcus* isolates (**Fig. 2B**). Conversely, some bacterial isolates (*Serratia marcesesens 855, Proteus mirabilis, Lactobacillus casei*) were bound by BrmIgA from few donors (**Fig. 2B**). The heterogeneity of anti-bacterial IgA reactivity was also demonstrated by comparison of BrmIgA responses to related isolates of *E. coli* where the magnitude of response to each isolate of *E. coli* was highly individual to the donor and could vary more than 5-fold (**Fig. 2C**). Species and isolate level heterogeneity was also evident in responses against *Staphylococcus, Serratia, Klebsiella* and *Enterococcus* (**Fig. 2A-B**). Despite this evident heterogeneity, we wanted to measure whether any of the anti-bacterial IgA responses were correlated, such that response to one bacterium would be predictive of another. To test this possibility, we used correlation network analyses to identify statistically significant pairwise relationships between different bacteria isolates (Ackerman et al., 2018; Suscovich et al., 2020). These analyses focus on the most significant pairwise correlations of IgA binding profiles across the different bacterial isolates and revealed substantial interconnection and correlated responses specific to *Enterobacteriaceae* isolates (**Fig. 2D**). This finding is consistent with a previous discovery of a high degree of *Enterobacteriaceae* cross-reactivity in blood-derived human IgA clones, due to reactivity to shared surface molecules, such as lipopolysaccharide (Rollenske et al., 2018). To gain a more comprehensive understanding of global relationships in IgA binding profiles across bacterial isolates, we visualized all pairwise correlations in a heatmap (**Fig. 2E**). The correlation heatmap showed two clear blocks – one composed entirely of *Enterobacteriaceae* involving highly correlated profiles, and the other involving relatively uncorrelated profiles. There were no strong correlations discovered amongst Gram Positive bacteria, even when comparing isolates of the same bacterial species (**Fig. 2D-E**). Taken together, our data indicates that even though anti-*Enterobacteriaceae* IgA responses are common and broad, the breast milk-derived anti-bacterial BrmIgA response is quite heterogeneous from person to person. Heterogeneity in anti-bacterial IgA may be important to newborns where IgA binding (or lack thereof) to infant intestinal bacteria can regulate bacterial colonization.

**Figure 2.**
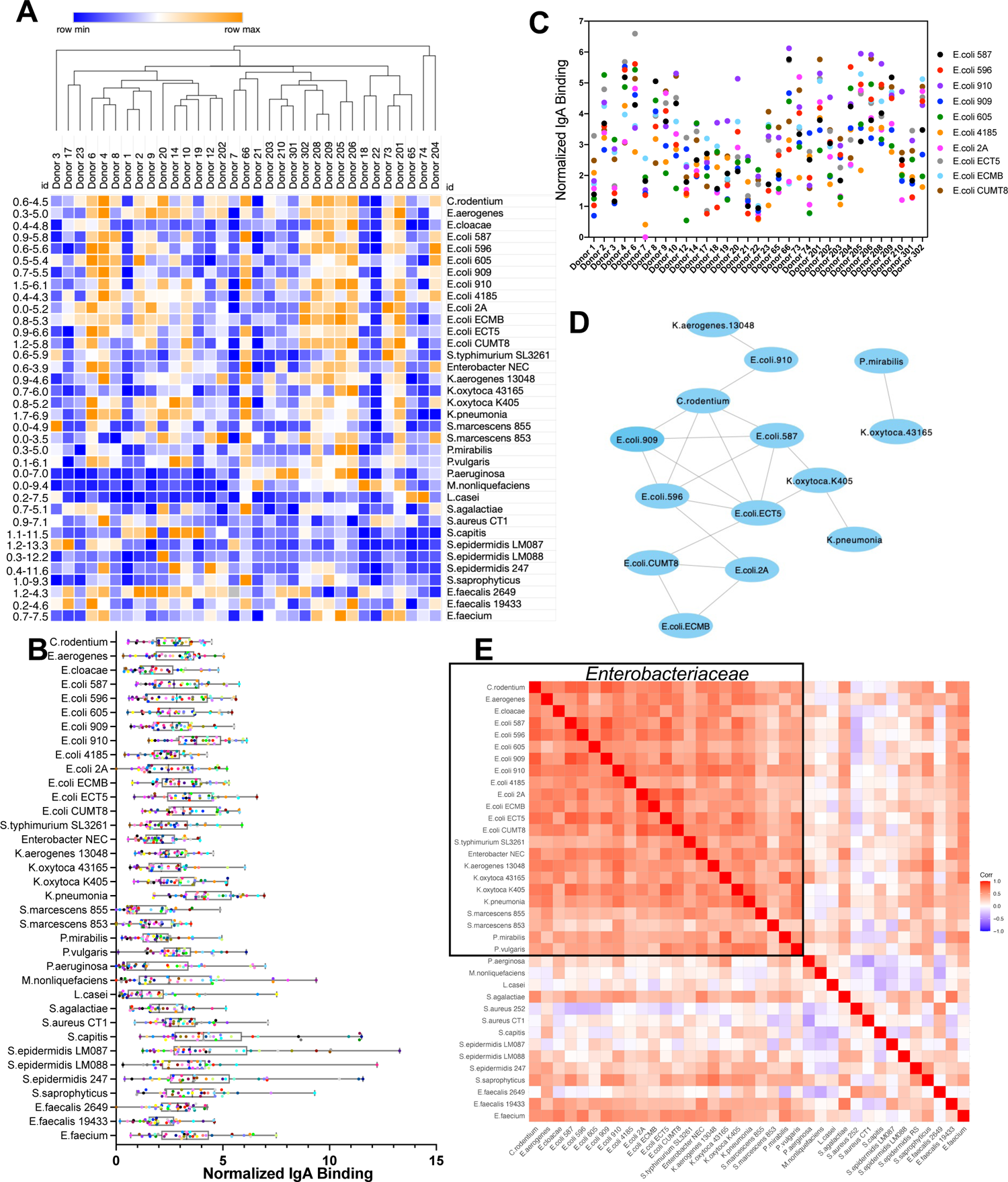
Heterogeneity in the anti-bacterial reactivity of breast milk-derived IgA. Donor milk samples (term infants; >37 weeks gestational age) were analyzed with our flow cytometric array (**1A). A)** Heat map of normalized anti-bacterial IgA binding affinity of different donors. Hierarchical clustering (Spearman). The range of the normalized values across each row is indicated on the left hand column. **B)** Scatter graph showing the normalized anti-bacterial IgA binding values for each donor (each color represents a different donor). **C)** Scatter graph of the normalized BrmIgA binding to different isolates of *E. coli* separated according to donors selected from the analysis in (**2A**). **D-E)** A correlation network analysis was performed to describe which anti-bacterial IgA responses were predictive. **D)** Network diagram indicating significantly correlated anti-bacterial IgA responses. **E)** Heat map indicating the level of correlation between different bacteria in our array. Black box drawn around *Enterobacteriaceae* family taxa.

### Heterogeneity in the breast milk-derived anti-bacterial IgA reactivity from donors who delivered preterm infants

It is not known when during pregnancy-induced mammary gland (MG) development that B cells traffic from the intestine to the MG. In mice it predominantly occurs late in gestation or even after delivery, whereas in pigs it occurs maximally in the 2^nd^ trimester (Langel et al., 2019; Roux et al., 1977). If B cell traffic to the MG during the 3^rd^ trimester is required for optimal breast milk IgA secretion in humans, preterm delivery, which often occurs at the transition between the 2^nd^ and 3^rd^ trimesters, may affect the level and specificity of breast milk-derived IgA. Comparison of the concentration of IgA from milk samples derived from preterm mothers (Gestational age 24-35 weeks) and term (>37 weeks) samples revealed no significant difference (**Fig. 3A**). Further, BrmIgA isolated from preterm milk samples phenocopied term milk samples with regard to heterogeneity within the anti-bacterial reactivity (**Fig. 3B-C**). Finally, PCA could not separate term and preterm samples on the basis of the anti-bacterial binding reactivity, indicating that, by the metrics of IgA concentration in the milk and anti-bacterial binding, preterm and term BrmIgA are effectively indistinguishable (**Fig. 3D**).

**Figure 3.**
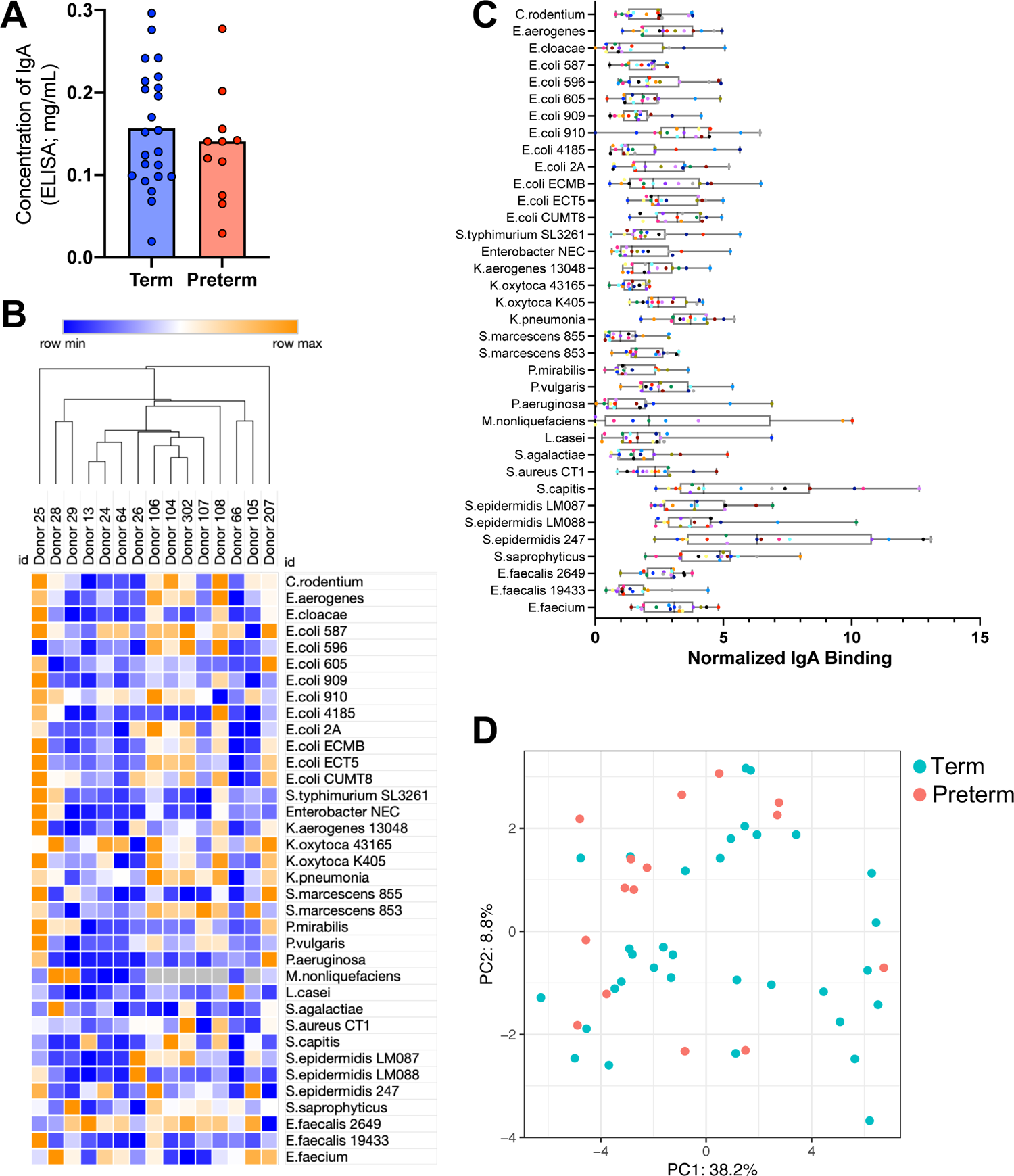
Heterogeneity in the breast milk-derived anti-bacterial IgA reactivity from donors who delivered preterm infants. Donor milk samples (preterm infants; 24-35 weeks gestational age) were analyzed with our flow cytometric array (**1A). A)** Bar graph showing the concentration of IgA purified from donor milk samples from mothers of term and preterm infants (ELISA). **B)** Heat map of normalized anti-bacterial binding affinity of different preterm donors. (Spearman). Samples where no data was collected due to insufficient bacteria in the well are colored grey. **C)** Scatter graph showing the normalized anti-bacterial IgA binding values for each preterm donor (each color represents a different donor). **D)** Principal Component Analysis (PCA) comparing aggregate anti-bacterial IgA binding between preterm and term samples.

### Temporal stability of anti-bacterial maternal IgA reactivity within one childbirth/infant

The concentration of all proteins, including BrmIgA, is highest in colostrum and then recedes to a stable point after the transition into mature milk, but whether this shift is associated with changes in the anti-bacterial reactivity of BrmIgA is not known. For example, whether B cells traffic in and out of the mammary gland during lactation, thus changing BrmIgA reactivity, is not well understood. We tested the stability of the anti-bacterial BrmIgA reactivity of various milk donors throughout their lactation periods, testing samples from each of the stages (colostrum, transitional, mature). All samples were normalized for the input concentration of IgA, which is critical to account for the increased level of IgA in colostrum. Hierarchical clustering of the longitudinal samples from the seven donors revealed that in general, samples captured from the same donor over long periods of time generally clustered together, indicating that they resemble other samples from the same donor more than they resemble samples from another donor (**Fig. 4A**). Longitudinal comparison of the magnitude of the IgA response against each bacterial isolate also revealed no generalizable trend towards increased or decreased anti-bacterial IgA binding as samples transitioned from colostrum/ transitional milk to mature milk (**Fig. S2**). Graphical depiction of the magnitude of the response against each bacterial isolate also revealed ‘clustering’ of samples within donor groups (**Fig. 4B**; each donor is one color). Finally, PCA analysis demonstrated that the collections of samples from different donors generally formed distinct clusters (**Fig. 4C and D**). Similar to data captured from individual mothers of term and preterm infants (**Fig. 2 and 3**), longitudinal samples show significant heterogeneity between donors. Thus, while the anti-bacterial BrmIgA reactivity of each donor is distinct, within each mother, the anti-bacterial antibodies and perhaps mammary gland resident B cells are stable.

**Figure 4.**
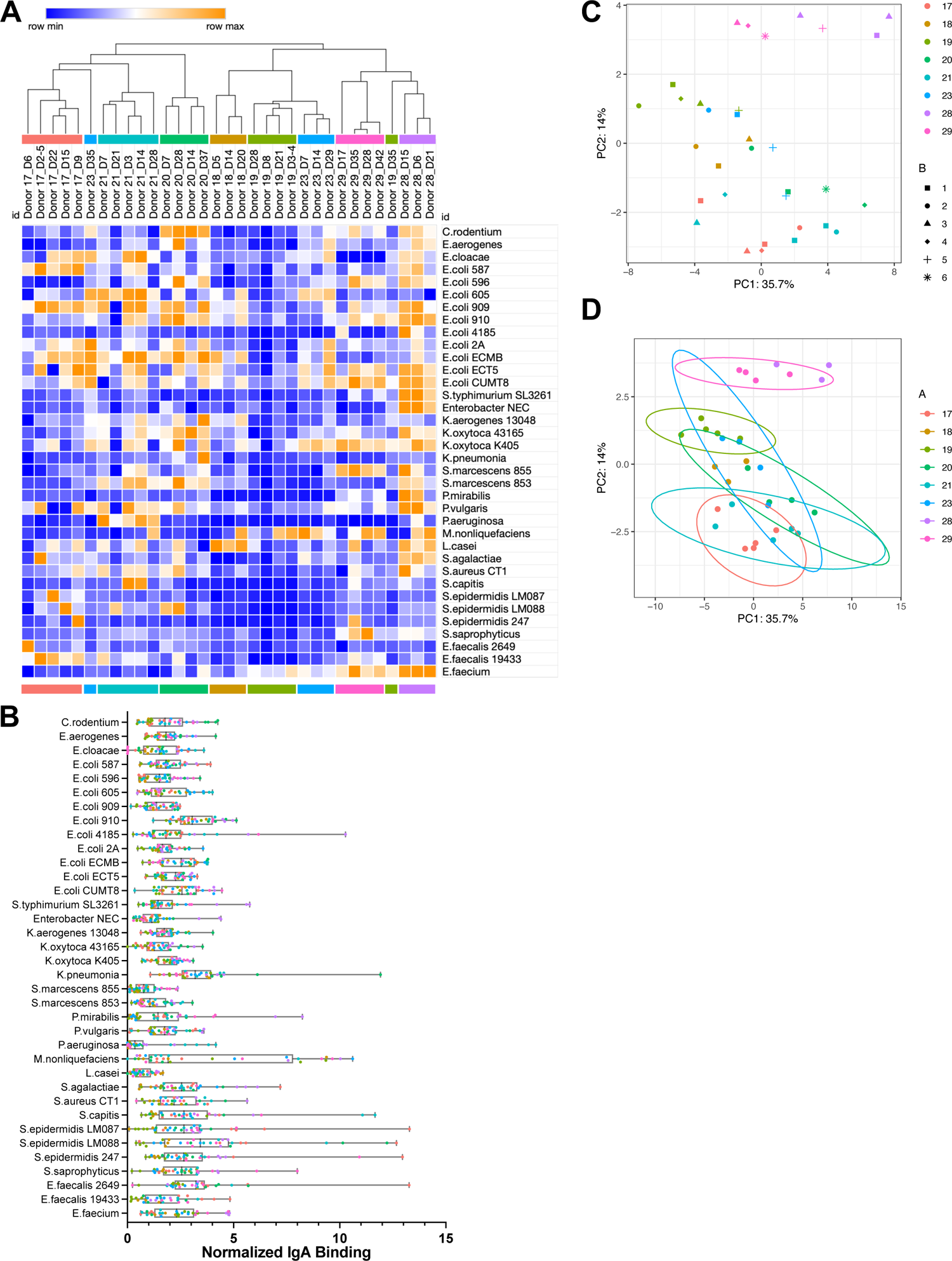
Temporal stability of anti-bacterial maternal IgA reactivity within one childbirth/infant. Multiple milk samples were collected from different donors over time and analyzed with our flow cytometric array (**1A**). **A)** Heat map of normalized anti-bacterial binding affinity of different donors. Hierarchical clustering (Spearman) of various donors is indicated by colored bars above and below the heatmap. Date of collection indicated on heatmap: D##, where the number is days post-delivery) **B)** Scatter graph showing the normalized anti-bacterial IgA binding values for each sample from longitudinally collected donors (each color represents a different donor; from **4A**). **C-D)** PCA of the aggregate anti-bacterial IgA binding of longitudinally collected samples. Each donor colored as in **4A**. **C)** PCA of individual longitudinally collected samples where symbols indicate the time of collection (week post delivery). **D)** PCA from **C** where ellipses indicate the maximum variance for each donor cluster along each axis. No ellipses are drawn for samples where fewer than four samples were available.

### Relative stability of the breast milk anti-bacterial IgA through siblings

During pregnancy, B cells are induced to traffic from the small intestine and Peyer’s Patches to the mammary gland (Lindner et al., 2015; Ramanan et al., 2020; Wilson and Butcher, 2004). In contrast to vaccine-specific B cells, microbiota-specific plasma B cells in the small intestine are believed to be replaced at a high rate and thus we hypothesized that the anti-bacterial reactivity of BrmIgA might shift substantially between sequential childbirths (Bemark et al., 2016; Hapfelmeier et al., 2010; Landsverk et al., 2017). To test this hypothesis, we acquired samples from a single donor over sequential infants and analyzed for changes in their anti-bacterial reactivity. Hierarchical clustering of these samples revealed that they clustered together and that samples captured after the 2^nd^ childbirth were more similar to the previous sample from the same individual than any other donor (**Fig. 5A**). PCA analysis confirmed the similarity of samples from sequential infants (**Fig. 5B**; comparison of the location of the same colored circles and triangles). There were individualized changes in IgA binding to different bacterial isolates between infants, but no generalizable bacterial isolate-specific trends were detected in the dataset between siblings (**Fig. 5A**). However, paired analysis of each multi-infant couplet comparing the mean change in anti-IgA binding across all of the isolates revealed that for the majority of donors anti-bacterial binding increased (6/10; upward pointing triangles) or stayed the same (2/10; circles) from the 1^st^ childbirth to subsequent childbirths (**Fig. 5C** and **Fig. S3**). Thus, we have observed that even between childbirths there is some stability in the anti-bacterial reactivity, implying either that B cells can permanently reside in the mammary gland outside of periods of lactation, or alternatively that the same or similar B cells are trafficking from mucosal sites during each pregnancy.

**Figure 5.**
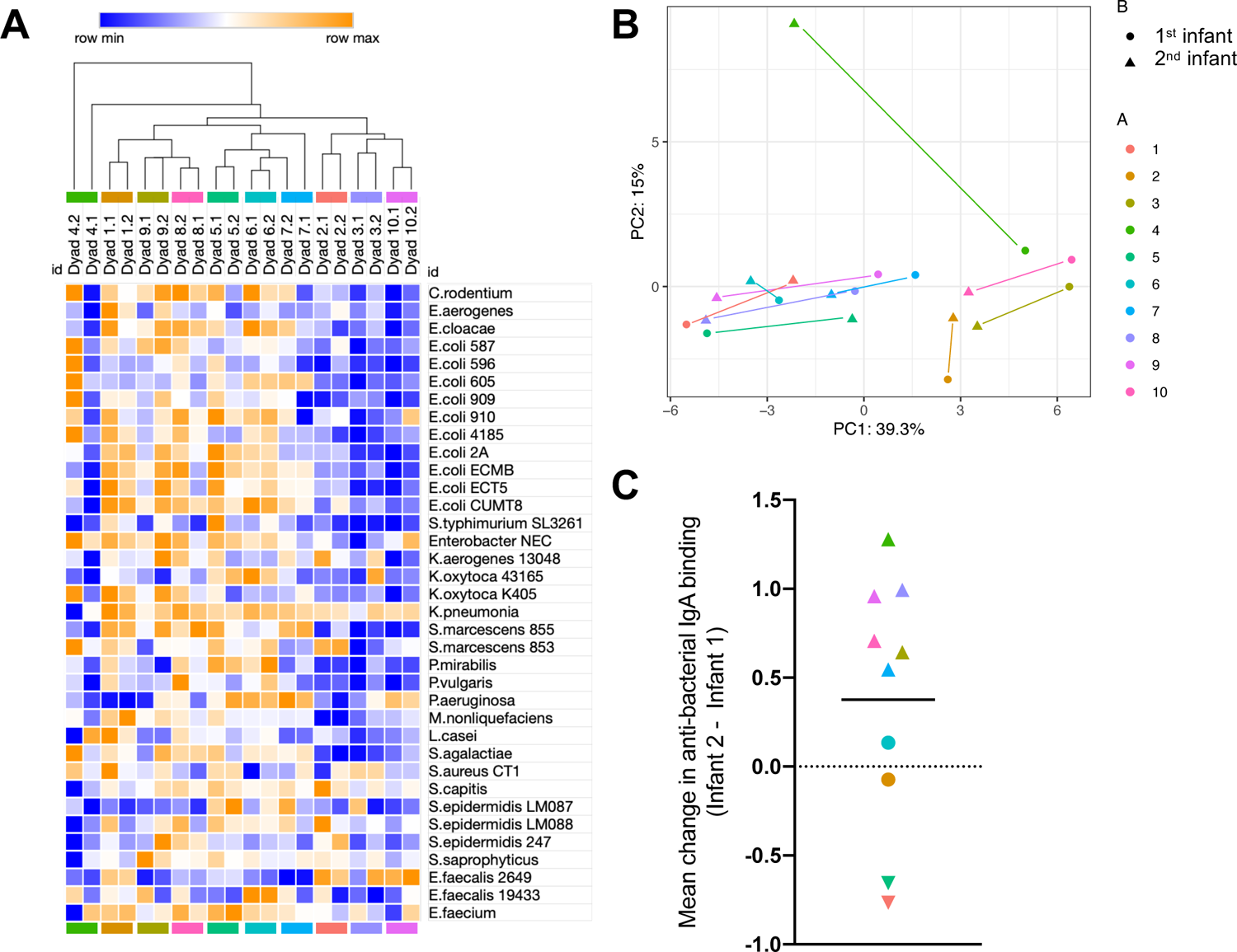
Stability of the breast milk-derived anti-bacterial IgA reactivity through sibling infants. Breast milk samples were collected from consecutive siblings and analyzed with our flow cytometric array (**1A**). **A)** Heat map of normalized anti-bacterial binding affinity of different donors. Hierarchical clustering (Spearman) of various donors is indicated by colored bars above and below the heatmap that correspond to each donor. **B)** PCA of aggregate anti-bacterial samples where each donor is displayed in a different color (from **5A**). The first sibling is indicated by a circle and the second sibling a triangle. Samples colored as in **5A. C)** Paired Student’s t-tests were calculated comparing the IgA binding of each donor between infant one and infant two for each bacterial taxon. The mean change ((Infant 2 – Infant 1; taxa 1) + (Infant 2 – Infant 1; taxa x))/37 (#of taxa) for each paired test was calculated and graphed. Significant increase in 2^nd^ infant = ‘up’ triangle; significant decrease in 2^nd^ infant = ‘down’ triangle; no statistical significance = circle. Colors are according to **5A**. See Supplemental Figure 3 for each Paired Student’s t test.

### Holder Pasteurization reduces the bacterial binding properties of breast milk-derived IgA

Increasingly, donor milk is being used as a substitute for Mother’s Own Milk (MOM) (Haiden and Ziegler, 2016). Donor milk has been shown to provide some of the benefits of MOM, including a reduction in the incidence of NEC, compared to formula-fed infants (Boyd et al., 2007; Canizo Vazquez et al., 2019; Miller et al., 2018; Quigley et al., 2018). To prevent the transfer of potentially pathogenic bacteria, donor milk is pasteurized by the Holder method (62.5°C for 30 minutes). An unfortunate consequence of Holder pasteurization is the denaturation of proteins and a reduction in the function of many of the immunological components of breast milk (Adhisivam et al., 2018). Secretory IgA is particularly stable, but it has been estimated that ∼13-62% of IgA is lost by Holder Pasteurization (Adhisivam et al., 2018; Lima et al., 2017; Peila et al., 2016). Here we split four donor samples in two and compared IgA concentrations and anti-bacterial IgA binding between raw control and Holder pasteurized samples. ELISA for the concentration of IgA before and after pasteurization revealed a 2-3-fold drop in the concentration of IgA, consistent with published literature (**Fig. 6A**). Critically, after normalizing for protein content between paired pasteurized and control samples we still detected an additional reduction in BrmIgA anti-bacterial binding responses to most isolates assayed (**Fig. 6B**). Thus, Holder pasteurization reduces both the amount and anti-bacterial binding capability of breast milk-derived IgA.

**Figure 6.**
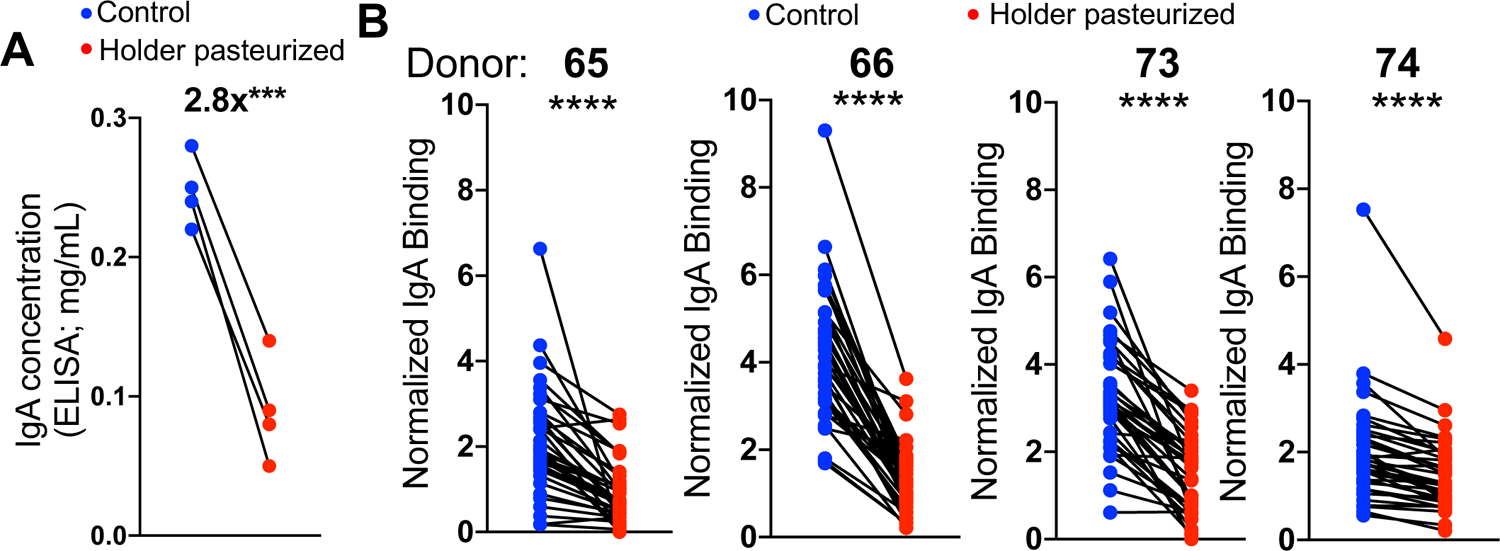
Holder pasteurization reduces the bacterial binding properties of breast milk-derived IgA. Breast milk samples from four donors were split into two where one half was pasteurized (62.5°C for 30 minutes) while the other untreated as a control. IgA was then isolated from both halves and analyzed on our flow cytometric array (**1A**). **A)** Paired Student’s t-test (***p<0.001) of the IgA concentration (mg/mL) of control (blue) and pasteurized samples (red), as measured by ELISA. **B)** Paired Student’s t-tests comparing control (blue) and Holder pasteurized (red) milk samples from the same donor. Each dot represents a different bacterial taxon. ****p<0.0001.

## Discussion

Here we demonstrate that the anti-bacterial reactivity of IgA in breast milk is heterogeneous between individuals but stable over time, both within one infant and over multiple childbirths. We did not find any appreciable difference in IgA content or functionality between preterm and term mothers. Additionally, we found that Holder pasteurization generally reduces the ability of breast milk-derived BrmIgA to bind bacteria, regardless of the identity of the bacteria.

A limitation of our study is the lack of obligate anaerobic bacteria within our flow cytometric array. We focused upon the facultative anaerobes that dominate the early colonization period of the infant because there is evidence that this is a critical time when BrmIgA is necessary to control microbiota colonization (Gopalakrishna et al., 2019; Mirpuri et al., 2014; Rognum et al., 1992), and that failure to control facultative anaerobes (*Enterobacteriaceae, Enterococcaceae, Streptococcaceae* etc.) is related to the development of NEC and other infant diseases (Flannery et al., 2021; Lin et al., 2022; Olm et al., 2019; Warner and Tarr, 2016). Obligate anaerobes are also problematic substrates for our flow cytometric array, which requires multiple staining and centrifugation steps difficult to perform in an anaerobic chamber and we are concerned that exposure to oxygen might kill or modify the bacteria leading to misleading results.

It is not surprising that each donor in our study possessed a distinct collection of anti-bacterial antibodies as this is likely the result of distinct life histories with regard to gastrointestinal infection and microbiota composition. Interestingly, we observed differences in the ability of individual donors to bind different isolates from the same species of bacteria. T cell-dependent IgA producing B cells are more likely to be targeted to bacterial surface proteins and less likely to be specific to repetitive structures on the bacteria’s surface. Thus, our findings support the hypothesis, derived from experiments in mice, that the majority of mammary gland resident IgA-producing B cells are the product of T cell-dependent activation (Bunker et al., 2017). We hypothesize that the specificity of milk-derived IgA is skewed towards heterogeneous surface proteins that differ between isolates of the same species, contributing to isolate level heterogeneity in IgA binding. Non-proteinaceous antigens (such as LPS) are also incredibly diverse in different bacterial isolates and are likely to contribute to the heterogeneity of breast milk-derived anti-bacterial reactivity. Conversely using network analyses, we do see some evidence of correlation between breast milk-derived anti-bacterial IgA responses directed against various *Enterobacteriaceae* family bacteria. Perhaps *Enterobacteriaceae* share surface structures to a greater degree than other bacteria we tested in our array, increasing the likelihood of IgA cross-reactiviy (Rollenske et al., 2018). It is possible that if we expanded the numbers of Gram Positive bacteria in the array that we would see more evidence of cross-reactivity, but we should note that amongst six isolates of *Staphylococcaceae* we observed no correlation in antibody responses.

In contrast to the heterogeneity that we observed between donors, we observed little heterogeneity in samples captured at different stages (from the same donor). This indicates that B cells may become established in the developing mammary gland and do not turn over to a substantial degree over the course of one infant. Indeed, the same B cell clones can be identified in breast milk samples over multiple timepoints (Bondt et al., 2021). This is important and underscores a key limitation of vertical antibody transmission into infants, which is that the maternal IgA response is physically separate from the target of its protective effect (infant’s intestine) and thus it does not respond to either bacterial or viral colonization of the infant. This is highly relevant to diseases common to preterm infants such as NEC and sepsis, where IgA present in breast milk may help prevent invasion by the nascent microbiota (Gopalakrishna et al., 2019). However, our results indicate that in some circumstances BrmIgA might not bind all infant intestinal bacteria and these ‘holes’ in anti-bacterial reactivity would persist throughout the breast feeding period, allowing unbound bacteria to proliferate and colonize more effectively. Previously, we observed a drop in IgA binding of *Enterobacteriaceae* that proceed the development of NEC (Gopalakrishna et al., 2019) and our new data implies that this observation is due a shift in the microbiota to escape maternal IgA and not change in the anti-bacterial IgA reactivity of the milk. Thus, for particularly at-risk preterm infants, it may be helpful to supplement breast milk with IgA known to bind the bacteria best associated to diseases like NEC.

We also observed that the anti-bacterial reactivity of BrmIgA was stable within one donor over multiple childbirths. This is somewhat surprising because the microbiota-specific B cells that populate the mammary gland traffic from the intestine and IgA producing B cells in the intestine are believed to turn over at a high rate (Hapfelmeier et al., 2010). Therefore, each pregnancy should lead to the deposition of new B cells and shifts in the anti-bacterial reactivity of BrmIgA. Our results demonstrated that BrmIgA reactivity from samples collected from one donor over several years and different infants looked more similar to each other than to any other donor, implying that either intestinal IgA-producing B cells are more stable than previously thought (Bemark et al., 2016), or that once established in the breast tissue, B cells can remain, even after lactation has been completed. In support of this idea, the majority of donors saw their responses either stay the same or improve in subsequent pregnancies. Whether B cells reside in mammary glands outside of the period of lactation and can be re-activated upon a subsequent pregnancy is testable in rodent models.

Feeding preterm infants human milk is well described to reduce the incidence of NEC compared to infant formula. Often for preterm infants, the mother’s milk production is insufficient and thus it is becoming more common to supplement the infant diet with pasteurized donor milk. Whether donor milk is as effective as Mother’s Own Milk (MOM) for protecting against NEC has not been conclusively determined (Quigley et al., 2018). Here we demonstrate that pasteurization reduces the both the amount of IgA in breast milk and the ability of BrmIgA to bind bacteria. Thus, if IgA is important for the effectiveness of donor milk in reducing NEC, one might suspect that donor milk would be less effective. Holder pasteurization also negatively affects other antibacterial proteins such as Lactoferrin, which could also reduce donor milk’s effectiveness (He et al., 2018; Pammi and Suresh, 2017). However, there are mitigating factors that might lessen the effects of pasteurization. First, donor milk provided to Neonatal Intensive Care Units is often a mixture of multiple donors, which almost certainly broadens the anti-bacterial reactivity, which could be beneficial. Second, we don’t actually know what the minimum functional amount of IgA binding to bacteria that is required to modulate intestinal colonization, partially because we do not fully understand the mechanism by which BrmIgA functions (Hand and Reboldi, 2021; Pabst and Slack, 2020; Yang and Palm, 2020). Finally, Human Milk Oligosaccharides, which shift the neonatal microbiota by increasing *Bifidobacteria,* are almost completely unaffected by pasteurization and very likely contribute to both preventing NEC and promoting a healthy infant microbiota (Bode, 2018).

Taken altogether we have demonstrated that there is substantial heterogeneity in the anti-bacterial reactivity of breast milk-derived IgA. We contend that this knowledge will serve as an important starting point for future studies on how binding by BrmIgA (or lack thereof) of newly colonizing bacteria shapes their ability to invade the infant intestine.

## Materials and Methods

### Study Design

#### Research objectives

Our objective was to identify the heterogeneity (or lack thereof) of breast milk derived IgA in response to common bacteria that colonize infants early after birth (in particular preterm infants).

#### Research subjects

De-identified milk donors from the Mid Atlantic Mother’s Milk bank (Pittsburgh, PA) and or Mommy’s Milk Human Milk Research Biorepository (San Diego, CA).

#### Experimental Design

We analyzed the anti-bacterial IgA reactivity with a custom bacterial flow cytometric array which we designed in our lab laboratory specifically for this purpose and is described in detail in both Figure 1 and below in the methods section. No randomization or blinding was used for this study.

#### Sample size

Since this was a discovery project and we really did not know the level of heterogeneity present within the breast milk-derived anti-bacterial IgA reactivity we did not perform a power analysis.

#### Data inclusion and exclusion

All samples that we acquired were analyzed, except samples where the IgA concentration was too low. In some cases, specific wells were omitted from our analysis if the number of bacteria in the well was insufficient for analysis (mostly *M. nonliquefaciens*). No outliers were excluded. All acquired data is included in our analyses.

#### Replicates

Samples were processed and analyzed over many weeks and consistent flow cytometric measurements (though heterogeneous between samples) were an important internal control that was continuously assessed. During the development of our methodology we repeated IgA/bacterial binding assays on consecutive days with the same milk-derived IgA samples and bacterial isolates to confirm that the staining was repeatable.

### Samples and Protocols

#### Human Donor Milk Samples

The human study protocol was deemed ‘Not Human Research’ by the Institutional Review Board (Protocol number PRO19110221) of the University of Pittsburgh. The majority of the de-identified donor maternal milk was acquired from the Mid-Atlantic Mothers Milk Bank DBA Human Milk Science Institute and Biobank of Pittsburgh, Pennsylvania. We acquired de-identified maternal milk collected over multiple childbirths (dyads) from Mommy’s Milk Human Milk Research Biorepository of San Diego, California. All donor milk samples were stored at −80° C.

#### Donor metadata

| Cohort | Term (PA) | Preterm (PA) | Longitudinal (PA) | Two infants (CA) | Pasteurization |
| --- | --- | --- | --- | --- | --- |

|  |  |  |  |  | **(PA)** |
| --- | --- | --- | --- | --- | --- |
| **Donor age (Years)** | 32.5+/-4.2 (29-40)<br>(unknown for 11 donors) | 32.5+/-4.5 (19-38) | 33.7+/-4.3 (29-40) (unknown for 1 donor) | 31.7+/-3.9 (22-39)<br>(includes age at 1<sup>st</sup> and 2<sup>nd</sup> infant) | 30.5+/-8.3 (19-39) |
| **Est. Gest. age of infant at delivery (Weeks)** | 39.6+/-+1.1 (37-41) | 30.7+/-3.1 (25-35) | 37.9+/-3.4 (32-41) | 38.2+/-3.4 (25-42)<br>(includes both infants) | 36.5+/-4.4 (30-39) |
| **Time post-delivery of sample collection (Days)** | 74.9+/-79.6 (14-276) | 55.6+/-100.3 (4-330) | 18+/-11.2 (3-42) | 258.3+/-157.6 (44-688)<br>(includes both infants) | 7.5+/-1 (7-9) |
| **Race of donor** | 100% Caucasian | 93% Caucasian<br>7% Black | 100% Caucasian | 20% Caucasian<br>40% Black<br>40% Asian | 100% Caucasian |
| **Ethnicity of donor** | Unknown | Unknown | Unknown | 40% Hispanic | Unknown |

Errors (+/-) represent the Standard Deviation. Number in brackets indicate range.
Metadata was not collected from all donors, as indicated. (PA = samples from Pittsburgh, PA; CA = samples from San Diego, CA)

#### Immunoglobulin A Extraction

To extract the IgA from milk, the donor milk was thawed at 4°C and 2mL of the maternal milk was placed in a 2ml Eppendorf tube. To separate the whey protein from the fat, the maternal milk was centrifuged at 16,000g for 5 mins at 4°C. The fat formed a layer at the top of the tube and the cells at the bottom of the tube. The whey protein was separated from the fat by carefully pipetting and filtering through a 0.22μm syringe filter, followed by washing the 0.22μm syringe filter with 500μL of wash buffer [Phosphate Buffered Saline (PBS)]. The filtered sample was then passed through a gravity flow column containing peptide M agarose after equilibrating the column with PBS. The sample was allowed to completely enter the matrix. The columns were washed with 20ml of 1X PBS. The column was then eluted with 10 mL elution buffer (0.1 M glycine, pH 2-3). 10ml of 1 M Tris with pH of 7.5 was used to neutralize the solution. The 20 mL sample was concentrated using a protein concentrator, by centrifugation of the column at 3000g for 20 min at 4° C. The concentrated sample was collected in 1.5 mL Eppendorf tubes and stored at −80° C.

#### Immunoglobulin A Quantification

Prior to running IgA samples on our array protein content was estimated via measurement on a Nanodrop UV Spectrophotometer. The concentration of IgA in each sample was measured by ELISA (Abcam) according to the manufacturer’s directions.

#### Protein separation and detection

Various fractions of either protein (pre and post-Peptide M column) were loaded onto a gradient acrylamide gel (4-15 %) and separated by LDS-page electrophoresis prior to staining with Coomassie Blue stain. Alternatively, proteins separated by weight were transferred onto nitrocellulose membranes and identified by Western Blotting. Membrane blocking and primary antibody staining was performed in TBS-Tween (X%) with the addition of powdered milk (anti-IgA Heavy Chain 1:10,000, Abcam; anti-Light Chain (kappa) 1:1000, Abcam; anti-J chain 1:500, ThermoFisher; anti-Secretory Factor 1:400, Abcam)

#### Bacterial Cultures and Flow cytometric Array development

We identified 13 genera commonly found within preterm infants and identified strains within the University of Pittsburgh community and ATCC collection that would be representative of the preterm infant microbiota. Bacteria were grown according to guidelines provided by ATCC or the providing investigator (see chart below), approximately 18-42 hours. The bacteria were diluted two-fold (1:2) to measure OD. 1mL of bacterial stock was then added to 1.5 mL eppendorf tube. The Eppendorf tube was centrifuged at 8,000g for 5 minutes and washed with 1 mL sterile 1X PBS twice. The supernatant was removed and re-suspended with 1 mL of sterile 1X PBS. The bacteria was then diluted to make the final concentration of 8 x 10^7^/mL Colony Forming Units (CFU). To preserve the integrity of the bacteria during freezing process 100μL of glycerol was added to the dilution (1:10). 27μL of the bacteria and glycerol mixture was then added to 2 wells each in a 96-well U-bottom plate, as experiment and control. The plates (containing 36 samples) plates are then stored at −80° C.

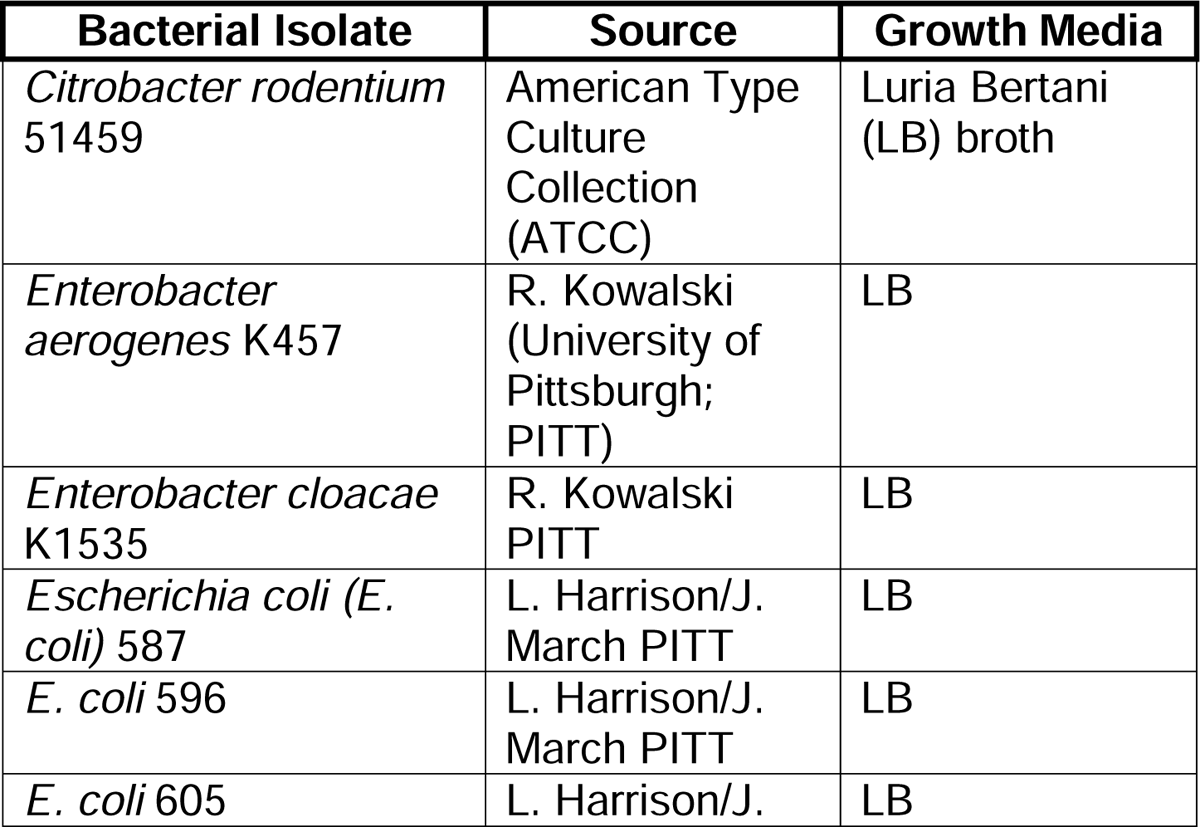
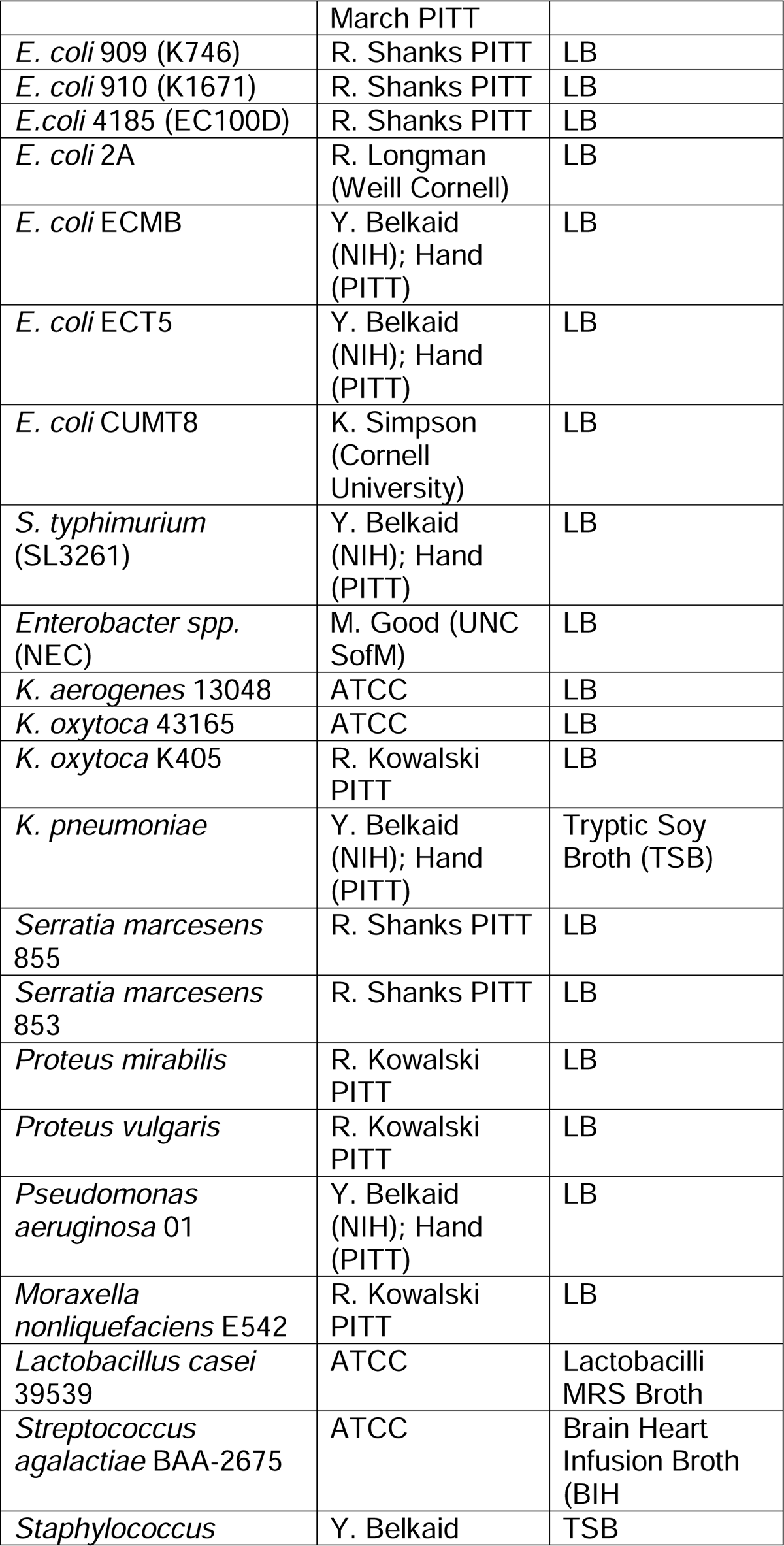
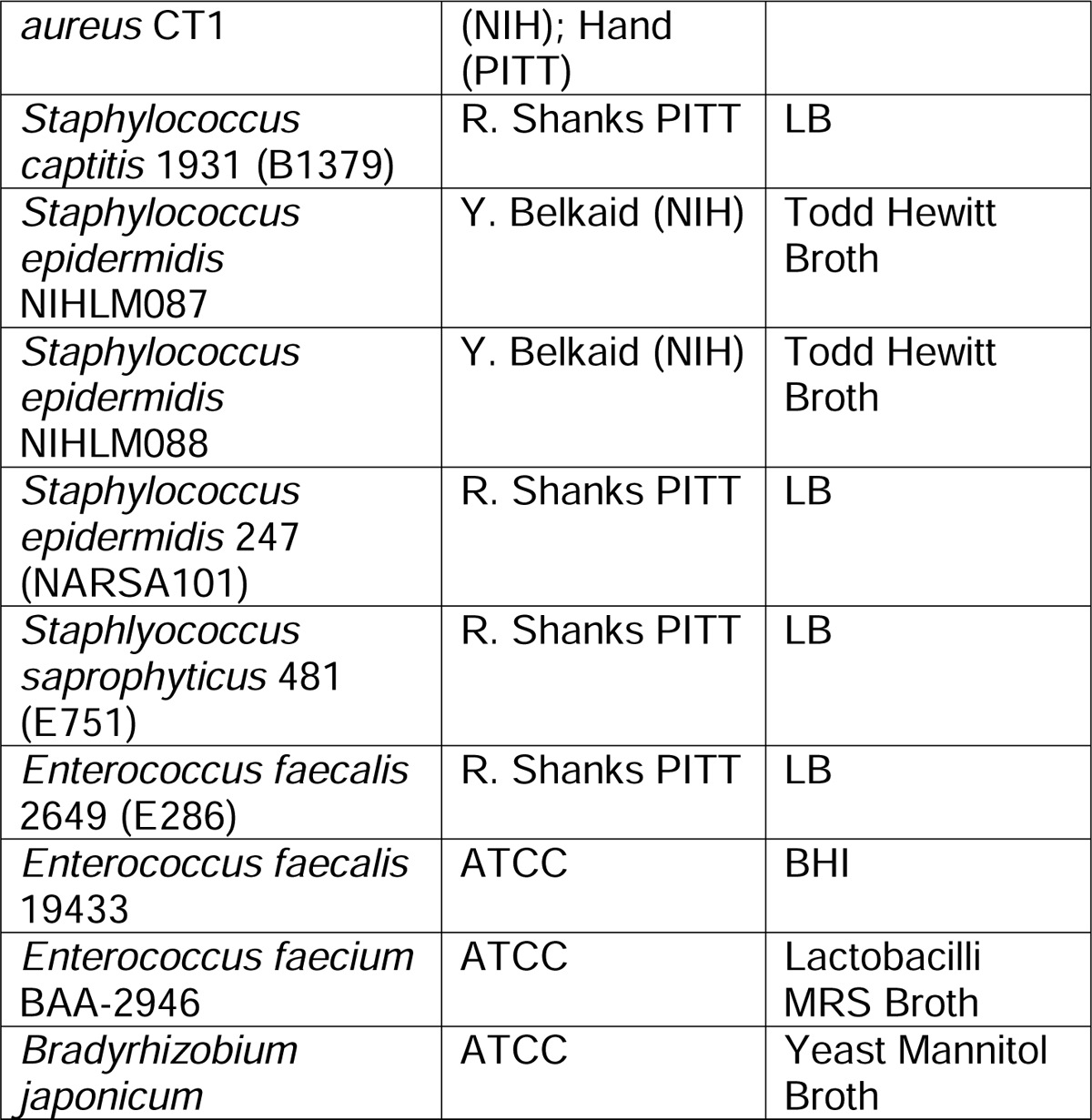

#### Bacterial Flow Assay

The bacterial plates, stored at −80°C were thawed at room temperature and washed twice (Swinging bucket centrifuge: 4,000 RPM for 5 minutes) with 200μL wash buffer [0.5% Bovine Serum Albumin (Sigma) in PBS-filtered through a 2.2μm filter]. The concentrated IgA from breast milk samples was thawed at 4°C and normalized to 0.1mg/ml by diluting the sample with sterile PBS. 25μL of the normalized IgA with 25μL of sterile 1X PBS was added to all the bacteria in the experimental wells. For controls, 50μL of sterile PBS was added. The plate was incubated for one hour in the dark on ice. After incubation, the plate was washed twice with 200μL wash buffer (4,000 RPM for 5 minutes). All the wells in the 96-well plate were then stained with 50μL secondary antibody staining mixture of Syto BC [(Green Fluorescent nuclear acid stain, Life Technologies (1:400)], APC Anti-Human IgA [Anti-Human IgA APC (Miltenyi Biotec clone REA1014, (1:50)], and blocking buffer of Normal Mouse Serum [ThermoFisher (1:10)]. The stained samples were incubated in the dark for an hour on ice. Samples were then washed three times with 200μL of wash buffer before flow cytometry analysis on the LSRFortessa-BD Biosciences.

For every donor we ran a separate plate that was stained only with the Syto BC/ APC anti-human IgA mix. These control samples are used as background fluorescence controls to establish positive binding signals and normalize samples collected on different days.

## Quantification and Statistical Analysis

### Flow Cytometry

All the data from flow cytometry was collected on a LSR Fortessa flow cytometer from BD Biosciences. The raw data was analyzed through the software FlowJo V10.4.2 (FlowJo, OR, USA). All samples were normalized according to the following formula: Log2 [(Geometric mean fluorescence intensity (gMFI) breast milk-derived IgA stained sample/ (gMFI of bacteria stained only with anti-human IgA APC antibody). Negative values (where the control has greater fluorescence than stained) are set to zero.

### Principal Component Analysis

Principal Component Analysis (PCA) plots were made using available R packages (ggplot2) and displays similarities in the percent binding of each donor sample to each bacterial taxon. Confidence ellipses demonstrate distinct groups based on multivariate t distribution.

### Correlation Network Analyses

We computed and visualized all pairwise Pearson correlations (of IgA binding profiles across bacterial isolates) in a heatmap using R. Significant correlations were defined using an effect size threshold of |r|>0.7 and an FDR (*P* value adjusted for multiple comparisons using Benjamini-Hochberg multiple testing correction) threshold of < 0.05. The significant correlations were visualized as a network using Cytoscape.

### Statistical Tests and analysis software

Heat maps were created using the MORPHEUS software tool (Broad Institute, Cambridge, MA). Hierarchical clustering by Spearman correlation. Samples collected over multiple infants were compared by either a standard or paired Student’s t-test (GraphPad PRISM 9).

## Summary of supplemental material

Three supplemental figures. Fig. S1 describes experiments related to the purification and quality control of IgA isolated from breast milk. Fig. S2 indicates the heterogeneity in BrmIgA anti-bacterial responses from longitudinally collected samples. Fig. S3 displays paired analysis of infant dyads who share a mother where we collected milk samples used to feed both infants.

## Acknowledgements

We would like to stress that we are only measuring the anti-bacterial IgA reactivity of breast milk samples and are not drawing conclusions on the best feeding option for any given infant. We would like to thank C. Verardi, D. O’Connor, K. Baumgartel and the entire staff of the Mid Atlantic Mother’s Milk Bank (Pittsburgh). We would also like to thank K. Bertrand and C. Chambers and the Mommy’s Milk Human Milk Research Biorepository of San Diego, California (UCSD). We would like to thank L. Harrison, R. Kowalski, J. Marsh, M. Morowitz and R. Shanks (Univ. of Pittsburgh, Pittsburgh, PA), R. Longman (Weill Cornell Medical School, New York, NY), Y. Belkaid (National Institutes of Health, Bethesda, MD), M. Good (University of North Carolina School of Medicine, Chapel Hill, NC) and K. Simpson (Cornell University, Ithaca, NY) for kindly providing bacterial strains and advice on strain selection. We would like to thank J. Michel and the Rangos Research Center Flow Cytometry core for assistance with running bacterial samples by flow cytometry. We would like to thank L. Cavacini for the anti-HIV IgA antibody. This work was supported by the UPMC Children’s Hospital of Pittsburgh/R.K. Mellon Institute for Pediatric Research, the NIH (R01DK120697 to T.W.H. and T32AI089443 to A.H.P.B.) and a March of Dimes Innovative Challenge Grant (T.W.H.). T.W.H. and K.P.G. have a patent for the technology used for the identification of anti-bacterial antibodies from breast milk samples (US17/341,272). T.W.H. has been engaged by Keller Postman LLC for his expertise on the components of breast milk that regulate intestinal colonization by bacteria and the development of Necrotizing Enterocolitis.

## Author contributions

C.B. Johnson-Hence, K. P. Gopalakrishna, K.E. Coffey, D. Bodkin and T.W. Hand conceptualized the project, developed the methodology and designed the experiments. C.B. Johnson-Hence, K. P. Gopalakrishna, K.E. Coffey, Y.A. Sosa, D. Bodkin, D.A. Abbott, A.T. Rai and J.T. Tometich performed the experiments. C.B. Johnson-Hence, D. Bodkin, A.H.P. Burr, S. Rahman, J. Das and T.W. Hand analyzed the data. C.B. Johnson-Hence and T. W. Hand wrote the manuscript.

## References

Ackerman, M.E., J. Das, S. Pittala, T. Broge, C. Linde, T.J. Suscovich, E.P. Brown, T. Bradley, H. Natarajan, S. Lin, J.K. Sassic, S. O’Keefe, N. Mehta, D. Goodman, M. Sips, J.A. Weiner, G.D. Tomaras, B.F. Haynes, D.A. Lauffenburger, C. Bailey-Kellogg, M. Roederer, and G. Alter. 2018. Route of immunization defines multiple mechanisms of vaccine-mediated protection against SIV. Nat Med 24:1590–1598.

Adhisivam, B., B. Vishnu Bhat, K. Rao, S.M. Kingsley, N. Plakkal, and C. Palanivel. 2018. Effect of Holder pasteurization on macronutrients and immunoglobulin profile of pooled donor human milk. J Matern Fetal Neonatal Med 1–4.

Bemark, M., H. Hazanov, A. Stromberg, R. Komban, J. Holmqvist, S. Koster, J. Mattsson, P. Sikora, R. Mehr, and N.Y. Lycke. 2016. Limited clonal relatedness between gut IgA plasma cells and memory B cells after oral immunization. Nat Commun 7:12698.

Bode, L. 2018. Human Milk Oligosaccharides in the Prevention of Necrotizing Enterocolitis: A Journey From in vitro and in vivo Models to Mother-Infant Cohort Studies. Front Pediatr 6:385.

Bondt, A., K.A. Dingess, M. Hoek, D.M.H. van Rijswijck, and A.J.R. Heck. 2021. A Direct MS-Based Approach to Profile Human Milk Secretory Immunoglobulin A (IgA1) Reveals Donor-Specific Clonal Repertoires With High Longitudinal Stability. Front Immunol 12:789748.

Boyd, C.A., M.A. Quigley, and P. Brocklehurst. 2007. Donor breast milk versus infant formula for preterm infants: systematic review and meta-analysis. Arch Dis Child Fetal Neonatal Ed 92:F169–175.

Bunker, J.J., S.A. Erickson, T.M. Flynn, C. Henry, J.C. Koval, M. Meisel, B. Jabri, D.A. Antonopoulos, P.C. Wilson, and A. Bendelac. 2017. Natural polyreactive IgA antibodies coat the intestinal microbiota. Science 358:

Canizo Vazquez, D., S. Salas Garcia, M. Izquierdo Renau, and I. Iglesias-Platas. 2019. Availability of Donor Milk for Very Preterm Infants Decreased the Risk of Necrotizing Enterocolitis without Adversely Impacting Growth or Rates of Breastfeeding. Nutrients 11:

Cortez, J., K. Makker, D.F. Kraemer, J. Neu, R. Sharma, and M.L. Hudak. 2018. Maternal milk feedings reduce sepsis, necrotizing enterocolitis and improve outcomes of premature infants. J Perinatol 38:71–74.

Dixon, D.L. 2015. The Role of Human Milk Immunomodulators in Protecting Against Viral Bronchiolitis and Development of Chronic Wheezing Illness. Children (Basel) 2:289–304.

Flannery, D.D., E.M. Edwards, K.M. Puopolo, and J.D. Horbar. 2021. Early-Onset Sepsis Among Very Preterm Infants. Pediatrics 148:

Gopalakrishna, K.P., and T.W. Hand. 2020. Influence of Maternal Milk on the Neonatal Intestinal Microbiome. Nutrients 12:

Gopalakrishna, K.P., B.R. Macadangdang, M.B. Rogers, J.T. Tometich, B.A. Firek, R. Baker, J. Ji, A.H.P. Burr, C. Ma, M. Good, M.J. Morowitz, and T.W. Hand. 2019. Maternal IgA protects against the development of necrotizing enterocolitis in preterm infants. Nat Med 25:1110–1115.

Haas, A., K. Zimmermann, F. Graw, E. Slack, P. Rusert, B. Ledergerber, W. Bossart, R. Weber, M.C. Thurnheer, M. Battegay, B. Hirschel, P. Vernazza, N. Patuto, A.J. Macpherson, H.F. Gunthard, and A. Oxenius. 2011. Systemic antibody responses to gut commensal bacteria during chronic HIV-1 infection. Gut 60:1506–1519.

Haiden, N., and E.E. Ziegler. 2016. Human Milk Banking. Ann Nutr Metab 69 Suppl 2:8–15.

Hand, T.W., and A. Reboldi. 2021. Production and Function of Immunoglobulin A. Annu Rev Immunol 39:695–718.

Hapfelmeier, S., M.A. Lawson, E. Slack, J.K. Kirundi, M. Stoel, M. Heikenwalder, J. Cahenzli, Y. Velykoredko, M.L. Balmer, K. Endt, M.B. Geuking, R. Curtiss, 3rd, K.D. McCoy, and A.J. Macpherson. 2010. Reversible microbial colonization of germ-free mice reveals the dynamics of IgA immune responses. Science 328:1705-1709.

He, Y.M., X. Li, M. Perego, Y. Nefedova, A.V. Kossenkov, E.A. Jensen, V. Kagan, Y.F. Liu, S.Y. Fu, Q.J. Ye, Y.H. Zhou, L. Wei, D.I. Gabrilovich, and J. Zhou. 2018. Transitory presence of myeloid-derived suppressor cells in neonates is critical for control of inflammation. Nat Med 24:224–231.

Johansen, F.E., and C.S. Kaetzel. 2011. Regulation of the polymeric immunoglobulin receptor and IgA transport: new advances in environmental factors that stimulate pIgR expression and its role in mucosal immunity. Mucosal Immunol 4:598–602.

Koch, M.A., G.L. Reiner, K.A. Lugo, L.S. Kreuk, A.G. Stanbery, E. Ansaldo, T.D. Seher, W.B. Ludington, and G.M. Barton. 2016. Maternal IgG and IgA Antibodies Dampen Mucosal T Helper Cell Responses in Early Life. Cell 165:827–841.

Landsverk, O.J., O. Snir, R.B. Casado, L. Richter, J.E. Mold, P. Reu, R. Horneland, V. Paulsen, S. Yaqub, E.M. Aandahl, O.M. Oyen, H.S. Thorarensen, M. Salehpour, G. Possnert, J. Frisen, L.M. Sollid, E.S. Baekkevold, and F.L. Jahnsen. 2017. Antibody-secreting plasma cells persist for decades in human intestine. J Exp Med 214:309–317.

Langel, S.N., F.C. Paim, M.A. Alhamo, A. Buckley, A. Van Geelen, K.M. Lager, A.N. Vlasova, and L.J. Saif. 2019. Stage of Gestation at Porcine Epidemic Diarrhea Virus Infection of Pregnant Swine Impacts Maternal Immunity and Lactogenic Immune Protection of Neonatal Suckling Piglets. Front Immunol 10:727.

Le Doare, K., B. Holder, A. Bassett, and P.S. Pannaraj. 2018. Mother’s Milk: A Purposeful Contribution to the Development of the Infant Microbiota and Immunity. Front Immunol 9:361.

Lima, H.K., M. Wagner-Gillespie, M.T. Perrin, and A.D. Fogleman. 2017. Bacteria and Bioactivity in Holder Pasteurized and Shelf-Stable Human Milk Products. Curr Dev Nutr 1:e001438.

Lin, Y.C., A. Salleb-Aouissi, and T.A. Hooven. 2022. Interpretable prediction of necrotizing enterocolitis from machine learning analysis of premature infant stool microbiota. BMC Bioinformatics 23:104.

Lindner, C., I. Thomsen, B. Wahl, M. Ugur, M.K. Sethi, M. Friedrichsen, A. Smoczek, S. Ott, U. Baumann, S. Suerbaum, S. Schreiber, A. Bleich, V. Gaboriau-Routhiau, N. Cerf-Bensussan, H. Hazanov, R. Mehr, P. Boysen, P. Rosenstiel, and O. Pabst. 2015. Diversification of memory B cells drives the continuous adaptation of secretory antibodies to gut microbiota. Nat Immunol 16:880–888.

Miller, J., E. Tonkin, R.A. Damarell, A.J. McPhee, M. Suganuma, H. Suganuma, P.F. Middleton, M. Makrides, and C.T. Collins. 2018. A Systematic Review and Meta-Analysis of Human Milk Feeding and Morbidity in Very Low Birth Weight Infants. Nutrients 10:

Mirpuri, J., M. Raetz, C.R. Sturge, C.L. Wilhelm, A. Benson, R.C. Savani, L.V. Hooper, and F. Yarovinsky. 2014. Proteobacteria-specific IgA regulates maturation of the intestinal microbiota. Gut Microbes 5:28–39.

Moor, K., J. Fadlallah, A. Toska, D. Sterlin, M.L. Balmer, A.J. Macpherson, G. Gorochov, M. Larsen, and E. Slack. 2016. Analysis of bacterial-surface-specific antibodies in body fluids using bacterial flow cytometry. Nat Protoc 11:1531–1553.

Neu, J., and W.A. Walker. 2011. Necrotizing enterocolitis. N Engl J Med 364:255–264.

Nino, D.F., C.P. Sodhi, and D.J. Hackam. 2016. Necrotizing enterocolitis: new insights into pathogenesis and mechanisms. Nat Rev Gastroenterol Hepatol 13:590–600.

Oddy, W.H. 2017. Breastfeeding, Childhood Asthma, and Allergic Disease. Ann Nutr Metab 70 Suppl 2:26–36.

Olm, M.R., N. Bhattacharya, A. Crits-Christoph, B.A. Firek, R. Baker, Y.S. Song, M.J. Morowitz, and J.F. Banfield. 2019. Necrotizing enterocolitis is preceded by increased gut bacterial replication, Klebsiella, and fimbriae-encoding bacteria. Sci Adv 5:eaax5727.

Pabst, O., and E. Slack. 2020. IgA and the intestinal microbiota: the importance of being specific. Mucosal Immunol 13:12–21.

Pammi, M., J. Cope, P.I. Tarr, B.B. Warner, A.L. Morrow, V. Mai, K.E. Gregory, J.S. Kroll, V. McMurtry, M.J. Ferris, L. Engstrand, H.E. Lilja, E.B. Hollister, J. Versalovic, and J. Neu. 2017. Intestinal dysbiosis in preterm infants preceding necrotizing enterocolitis: a systematic review and meta-analysis. Microbiome 5:31.

Pammi, M., and G. Suresh. 2017. Enteral lactoferrin supplementation for prevention of sepsis and necrotizing enterocolitis in preterm infants. Cochrane Database Syst Rev 6:CD007137.

Peila, C., G.E. Moro, E. Bertino, L. Cavallarin, M. Giribaldi, F. Giuliani, F. Cresi, and A. Coscia. 2016. The Effect of Holder Pasteurization on Nutrients and Biologically-Active Components in Donor Human Milk: A Review. Nutrients 8:

Planer, J.D., Y. Peng, A.L. Kau, L.V. Blanton, I.M. Ndao, P.I. Tarr, B.B. Warner, and J.I. Gordon. 2016. Development of the gut microbiota and mucosal IgA responses in twins and gnotobiotic mice. Nature 534:263–266.

Quigley, M., N.D. Embleton, and W. McGuire. 2018. Formula versus donor breast milk for feeding preterm or low birth weight infants. Cochrane Database Syst Rev 6:CD002971.

Ramanan, D., E. Sefik, S. Galvan-Pena, M. Wu, L. Yang, Z. Yang, A. Kostic, T.V. Golovkina, D.L. Kasper, D. Mathis, and C. Benoist. 2020. An Immunologic Mode of Multigenerational Transmission Governs a Gut Treg Setpoint. Cell 181:1276–1290 e1213.

Reyman, M., M.A. van Houten, D. van Baarle, A. Bosch, W.H. Man, M. Chu, K. Arp, R.L. Watson, E.A.M. Sanders, S. Fuentes, and D. Bogaert. 2019. Impact of delivery mode-associated gut microbiota dynamics on health in the first year of life. Nat Commun 10:4997.

Rogier, E.W., A.L. Frantz, M.E. Bruno, L. Wedlund, D.A. Cohen, A.J. Stromberg, and C.S. Kaetzel. 2014. Secretory antibodies in breast milk promote long-term intestinal homeostasis by regulating the gut microbiota and host gene expression. Proc Natl Acad Sci U S A 111:3074–3079.

Rognum, T.O., S. Thrane, L. Stoltenberg, A. Vege, and P. Brandtzaeg. 1992. Development of intestinal mucosal immunity in fetal life and the first postnatal months. Pediatr Res 32:145–149.

Rollenske, T., V. Szijarto, J. Lukasiewicz, L.M. Guachalla, K. Stojkovic, K. Hartl, L. Stulik, S. Kocher, F. Lasitschka, M. Al-Saeedi, J. Schroder-Braunstein, M. von Frankenberg, G. Gaebelein, P. Hoffmann, S. Klein, K. Heeg, E. Nagy, G. Nagy, and H. Wardemann. 2018. Cross-specificity of protective human antibodies against Klebsiella pneumoniae LPS O-antigen. Nat Immunol 19:617–624.

Roux, M.E., M. McWilliams, J.M. Phillips-Quagliata, P. Weisz-Carrington, and M.E. Lamm. 1977. Origin of IgA-secreting plasma cells in the mammary gland. J Exp Med 146:1311–1322.

Sandin, C., S. Linse, T. Areschoug, J.M. Woof, J. Reinholdt, and G. Lindahl. 2002. Isolation and detection of human IgA using a streptococcal IgA-binding peptide. J Immunol 169:1357–1364.

Slack, E., S. Hapfelmeier, B. Stecher, Y. Velykoredko, M. Stoel, M.A. Lawson, M.B. Geuking, B. Beutler, T.F. Tedder, W.D. Hardt, P. Bercik, E.F. Verdu, K.D. McCoy, and A.J. Macpherson. 2009. Innate and adaptive immunity cooperate flexibly to maintain host-microbiota mutualism. Science 325:617–620.

Sobti, J., G.P. Mathur, A. Gupta, and Who. 2002. WHO’s proposed global strategy for infant and young child feeding: a viewpoint. J Indian Med Assoc 100:502–504, 506.

Suscovich, T.J., J.K. Fallon, J. Das, A.R. Demas, J. Crain, C.H. Linde, A. Michell, H. Natarajan, C. Arevalo, T. Broge, T. Linnekin, V. Kulkarni, R. Lu, M.D. Slein, C. Luedemann, M. Marquette, S. March, J. Weiner, S. Gregory, M. Coccia, Y. Flores-Garcia, F. Zavala, M.E. Ackerman, E. Bergmann-Leitner, J. Hendriks, J. Sadoff, S. Dutta, S.N. Bhatia, D.A. Lauffenburger, E. Jongert, U. Wille-Reece, and G. Alter. 2020. Mapping functional humoral correlates of protection against malaria challenge following RTS,S/AS01 vaccination. Sci Transl Med 12:

Walker, W.A., and R.S. Iyengar. 2015. Breast milk, microbiota, and intestinal immune homeostasis. Pediatr Res 77:220–228.

Warner, B.B., and P.I. Tarr. 2016. Necrotizing enterocolitis and preterm infant gut bacteria. Semin Fetal Neonatal Med 21:394–399.

Wilson, E., and E.C. Butcher. 2004. CCL28 controls immunoglobulin (Ig)A plasma cell accumulation in the lactating mammary gland and IgA antibody transfer to the neonate. J Exp Med 200:805–809.

Yang, Y., and N.W. Palm. 2020. Immunoglobulin A and the microbiome. Curr Opin Microbiol 56:89–96.

Yu, X., M. Duval, C. Lewis, M.A. Gawron, R. Wang, M.R. Posner, and L.A. Cavacini. 2013. Impact of IgA constant domain on HIV-1 neutralizing function of monoclonal antibody F425A1g8. J Immunol 190:205–210.

Zhang, W., Q. Feng, C. Wang, X. Zeng, Y. Du, L. Lin, J. Wu, L. Fu, K. Yang, X. Xu, H. Xu, Y. Zhao, X. Li, U.H. Schoenauer, A. Stadlmayr, N.K. Saksena, H. Tilg, C. Datz, and X. Liu. 2017. Characterization of the B Cell Receptor Repertoire in the Intestinal Mucosa and of Tumor-Infiltrating Lymphocytes in Colorectal Adenoma and Carcinoma. J Immunol 198:3719–3728.

Zikan, J., J. Novotny, T.L. Trapane, M.E. Koshland, D.W. Urry, J.C. Bennett, and J. Mestecky. 1985. Secondary structure of the immunoglobulin J chain. Proc Natl Acad Sci U S A 82:5905–5909.

