## Supplemental Figures 1-3 for "Stability and heterogeneity in the anti-microbiota reactivity of human milk-derived Immunoglobulin A"

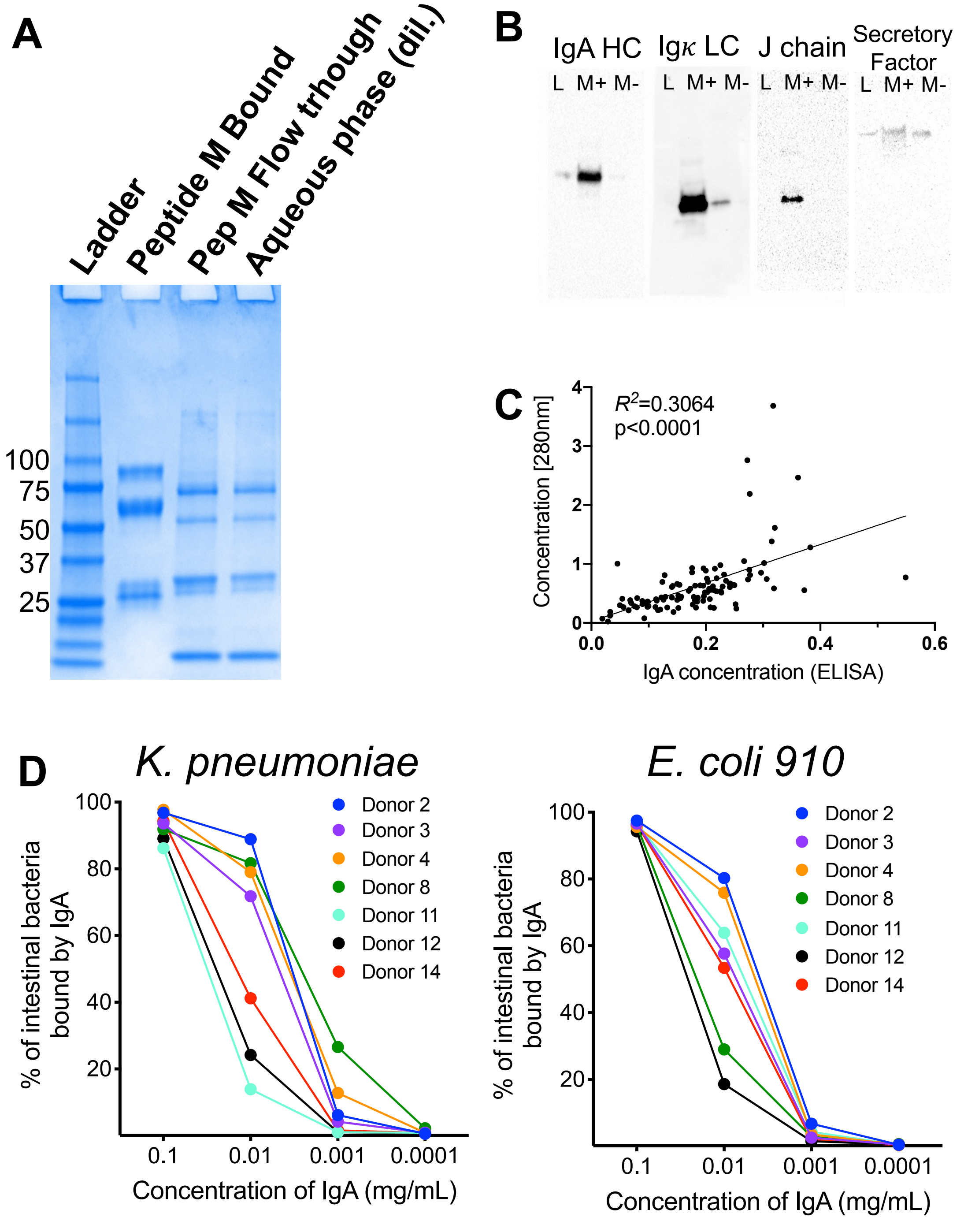


**Supplemental Figure 1 – Isolation of IgA from breast milk by Peptide M columns substantially enriches for secretory IgA.**

**A)** The soluble fraction of a breast milk sample was run over a Peptide M column and analyzed on an LDS-Page gel. Shown are the Peptide M bound fraction, the flow through and the aqueous phase prior to separation.

**B)** LDS-Page gels were prepared as in **SF1A,** transferred to nitrocellulose and blotted with antibodies to different proteins as described above the blot. L= ladder; M+ = Peptide M bound fraction and M- = flow through of Peptide M column.

**C**) Concordance of IgA estimations made my 280nm light absorbance and IgA ELISA.

**D)** 10-fold dilutions of Peptide M purified IgA fractions run over the flow cytometric array as described in 1A. Shown are two example bacteria.

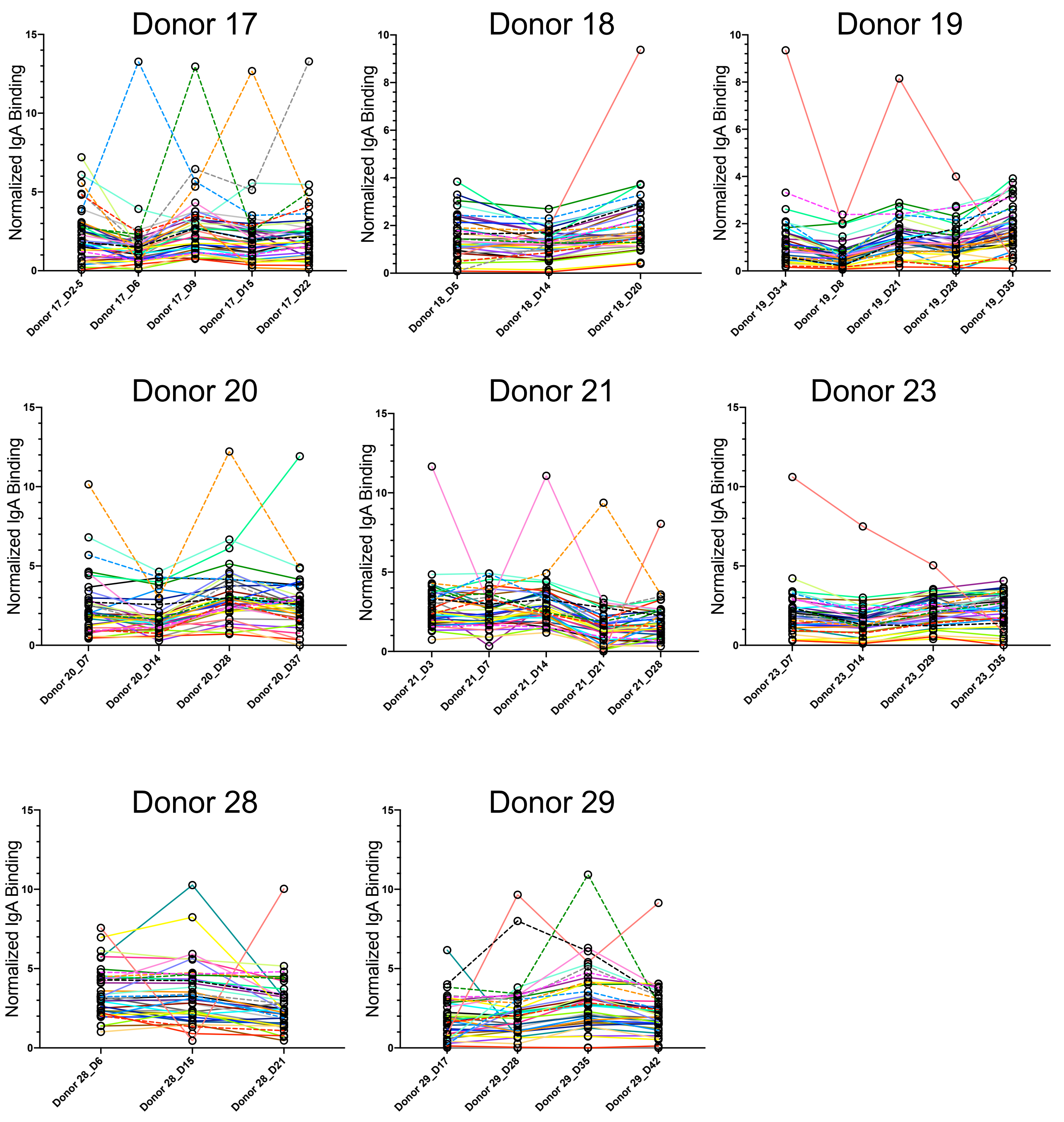


**Supplemental Figure 2 Longitudinal analysis of the anti-baterial IgA response from sequentially collected breast milk samples.**
Graph of normalized IgA binding to all bacterial types from longitudinal milk samples. Samples from each donor are in order left to right. Each donor is represented by an individual graph and each bacteria by a different colored line.

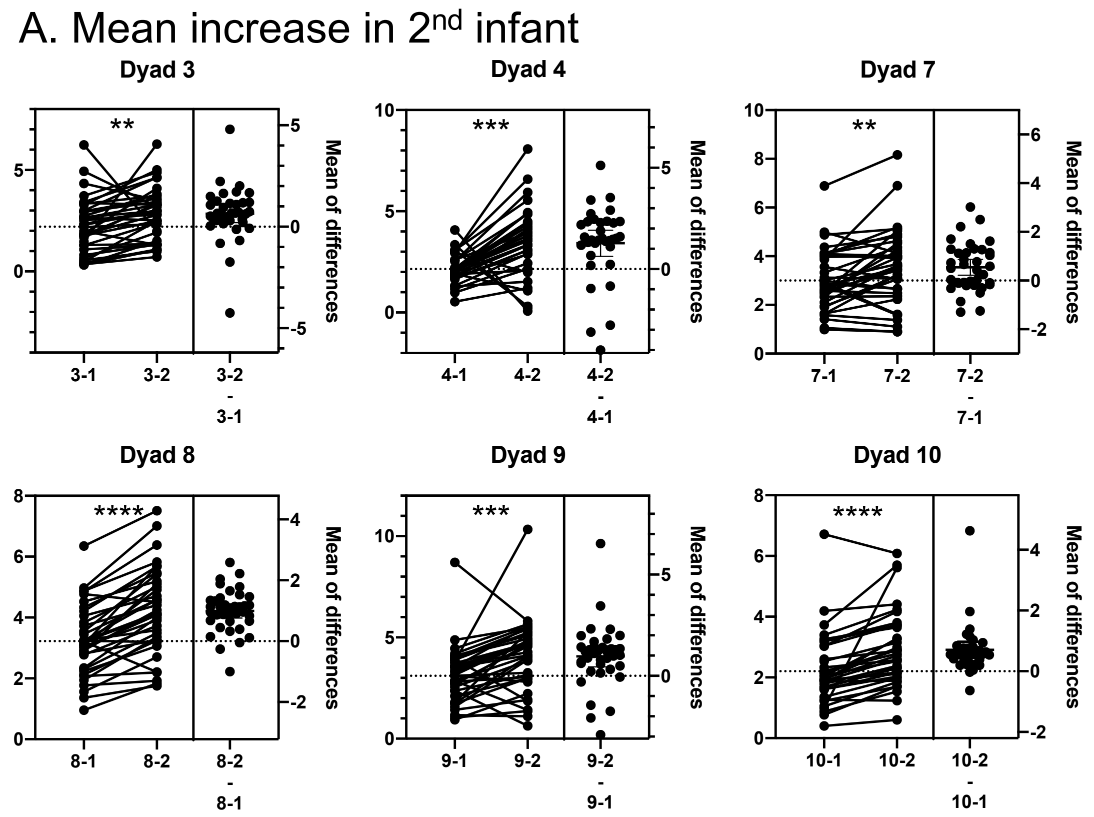


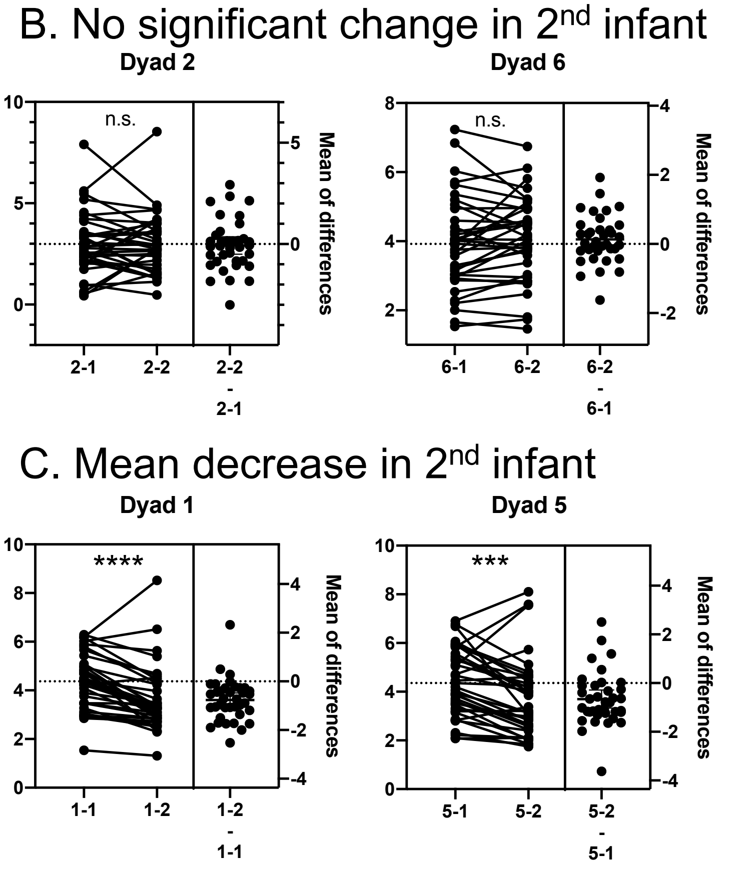


**Supplemental Figure 3 Detailed comparisons of the anti-bacterial IgA repertoire from one donor collected over two separate infants.** Paired Student's t-tests comparing the anti-bacterial IgA binding between breast milk collected from the first and second infant. Each dot represents a different bacterial taxon and each graph is a different donor (as indicated). The right-hand graph is the difference (norm. IgA binding infant 2- norm. IgA binding infant 1) for each taxon.
